# Startling acoustic stimuli hasten reflexive choice reaching tasks by strengthening, but not changing the timing of, express visuomotor responses

**DOI:** 10.1101/2024.07.01.601510

**Authors:** Vivian Weerdesteyn, Sarah L Kearsley, Aaron L Cecala, Ewan A Macpherson, Brian D. Corneil

## Abstract

Responding to an external stimulus takes ∼200 ms, but this can be shortened to as little as ∼120 ms with the additional presentation of a startling acoustic stimulus. This phenomenon is hypothesized to arise from the involuntary release of a prepared movement (a *StartReact* effect). However, a startling acoustic stimulus also expedites rapid mid-flight, reactive adjustments to unpredictably displaced targets which could not have been prepared in advance. We surmise that for such rapid visuomotor transformations, intersensory facilitation may occur between auditory signals arising from the startling acoustic stimulus and visual signals relayed along a fast subcortical network. To explore this, we examined how a startling acoustic stimulus shortens reaction times in a task that produces express visuomotor responses, which are brief bursts of muscle activity that arise from a fast tectoreticulospinal network. We measured express visuomotor responses on upper limb muscles in humans as they reached either toward or away from a stimulus in blocks of trials where movements could either be fully prepared or not, occasionally pairing stimulus presentation with a startling acoustic stimulus. The startling acoustic stimulus reliably produced larger but fixed-latency express visuomotor responses in a target-selective manner, and also shortened reaction times, which were equally short for prepared and unprepared movements. Our results provide insights into how a startling acoustic stimulus shortens the latency of reactive movements without full motor preparation. We propose that the reticular formation is the likely node for intersensory convergence during the most rapid transformations of vision into targeted reaching actions.

**KEY POINTS:**

- A startling acoustic stimulus (SAS) shortens reaction times by releasing fully prepared motor programs (the StartReact effect), but can also hasten responses in reflexive tasks without any movement preparation
- Here we measure the effect of a SAS on reaction times and upper limb muscle recruitment in a reflexive reaching task, focusing on express visuomotor responses that are evoked by visual target presentation and demarcate activity along a subcortical tectoreticulospinal pathway
- A SAS robustly increased the magnitude of express visuomotor responses without changing their timing, and this increase was tightly related to the subsequent reaction time even in the absence of motor preparation
- Our results attest to intersensory facilitation within the tectoreticulospinal pathway, which provides the shortest pathway mediating visuomotor transformations for reaching
- These results reconcile discrepant findings by emphasizing the importance of intersensory facilitation in SAS-induced hastening of reaction times in reflexive tasks

## Introduction

Initiation of voluntary movements to visual stimuli typically takes >200 ms. Yet, when a visual ‘Go’ stimulus is paired with a startling acoustic stimulus (SAS), reaction times (RTs) can be speeded up to a presumed ‘reactive’ mode of control with RTs of ∼80-120 ms (depending on whether EMG- or movement velocity-based readouts are reported; (Valls-Solé *et al*., 1995; Carlsen *et al*., 2004)). This shortening of RTs has been demonstrated in many *simple* reaction tasks involving single or multi-segmental arm and leg movements where movements can be fully prepared (for review see (Carlsen *et al*., 2012; Nonnekes *et al*., 2015)). However, the effect of a SAS is more nuanced in a *choice* reaction task which involves selecting between multiple responses. In such choice reaction tasks, a SAS typically does not generate ‘reactive’ RTs, and any large RT reductions in choice reaction tasks often come at a cost of increased errors or inaccuracies (Carlsen *et al*., 2004, 2009; Forgaard *et al*., 2011; Maslovat *et al*., 2012; Marinovic *et al*., 2017). The dependence on partial or full preparation prior to stimulus presentation has led to a mechanistic explanation of why a SAS shortens RTs, termed the *StartReact effect*, wherein the movement is involuntarily ‘released’ by the SAS (Valls-Solé *et al*., 1999; Carlsen *et al*., 2012; Carlsen & Maslovat, 2019).

However, there are reports of a SAS shortening RTs in choice reaction times, producing RTs just as fast to those observed for ‘prepared’ movements in simple reaction tasks, with neutral (Reynolds & Day, 2007; Queralt *et al*., 2008). Thus, under certain circumstances, a SAS facilitates rapid visuomotor transformations, even without a fully or partially prepared movement. Are such results also due a StartReact effect? One distinctive feature of these studies is that they both involved online movement corrections, in this case of the lower limb. Online movement corrections may represent a special class of reactive movements where visual input is directly mapped onto motor outputs via a fast subcortical network involving the tecto-reticulo-spinal system (Day & Lyon, 2000; Perfiliev *et al*., 2010; Kozak *et al*., 2019). Consistent with this, RTs of online corrections are very short even in the absence of a SAS (Soechting & Lacquaniti, 1983), such movements are initially directed invariably toward a visual stimulus (Day & Lyon, 2000), and their RTs do not follow Hick’s law as they remain fixed regardless of the number of possible alternatives (Reynolds & Day, 2012). Other reactive responses like express saccades are also invariably stimulus-driven and do not follow Hick’s law (Paré & Munoz, 1996), and are known to rely critically on the subcortical superior colliculus (Schiller *et al*., 1987; Edelman & Keller, 1996; Dorris *et al*., 1997). Rather than relying on the purported StartReact mechanism of involuntary movement release, could the hastening of RTs due to a SAS in reactive movements like online corrections arise instead from intersensory facilitation within the reticular formation between the SAS and visual signals relayed along a tecto-reticulo-spinal pathway? If so, then in some scenarios the effect of a SAS may not be to release a prepared motor program, but instead strengthen rather than expedite the rapid transformation of vision into action.

Recent work on intersensory facilitation across multiple sensory modalities suggests that a SAS may indeed strengthen the output of the fast visuomotor network (Glover & Baker, 2019). In the context of center-out visually-guided reaches from a stationary position in a choice reaction task, a SAS increased the mean magnitude of short-latency (∼80-120 ms) recruitment of upper limb muscles without drastically impacting its timing. Such recruitment may reflect what are termed express visuomotor responses (EVR; formerly termed stimulus-locked responses), which provide another measure of the output of the fast visuomotor network. The EVR is a brief increase/decrease in the target-selective recruitment of agonist/antagonist muscles that is relatively time-locked to the visual stimulus at a latency of ∼80-100 ms, and is spatially and temporally distinct from the longer-duration burst of muscle activity associated with the generation of the voluntary arm movement (Pruszynski *et al*., 2010; Wood *et al*., 2015; Gu *et al*., 2016; Atsma *et al*., 2018). Larger but fixed-latency EVRs precede shorter RTs, and there is compelling evidence that EVRs reflect tecto-reticulo-spinal processing (Pruszynski *et al*., 2010; Gu *et al*., 2016; Kozak *et al*., 2019; Contemori *et al*., 2021a, 2023; Kearsley *et al*., 2022; Selen *et al*., 2023; Billen *et al*., 2023). EVRs also precede and share many characteristics with the first phase of on-line corrections (Day & Lyon, 2000; Fautrelle *et al*., 2010; Gu *et al*., 2016; Kozak *et al*., 2019), consistent with the forces arising from the EVR serving to initiate on-line corrections. However, as Glover and Baker (2019) reported a generic enhancement of muscle recruitment with SAS across all target directions, it cannot be ruled out that this enhanced recruitment reflected generalized startle reflex-related potentiation, rather than target-selective facilitation of the EVR itself. If this were true, one would expect these SAS-enhanced EVRs to result in faster RTs for movements towards the body, but slower and with more frequent directional errors for those away from the body, due to the preferential recruitment of flexor muscles in the startle reflex (Brown *et al*., 1991b, 1991a). In contrast, in the event of intersensory facilitation of the fast visuomotor network itself, where the SAS presumably acts as an accessory stimulus to increase the excitation arising from the visual signal, RT shortening is expected in all directions in the absence of drastically increased errors. As the Glover and Baker (2019) study did not focus on movement initiation times, it is not known how the observed facilitation of the SAS on EVRs would compare across Choice and Simple reaction tasks, nor how trial-by-trial EVRs relate to the ensuing reactive RTs.

Here, we tested the hypothesis that simultaneous presentation of a SAS with a salient visual stimulus shortens the RTs of reactive reaching movements by strengthening the magnitude of EVRs without changing their latency. We used an emerging target paradigm that increases the generation of EVRs and reactive reaches, even in a choice reaction task on trials without a SAS (Kozak *et al*., 2020; Contemori *et al*., 2021b; Kozak & Corneil, 2021). In this task, EVRs are initiated when the subjects have not yet started to move, simplifying the quantification of muscle activity compared to an on-line correction task where the EVR evolves in concert with muscle recruitment associated with an ongoing movement. We also interleaved trials where subjects reached toward or away from the emerging stimulus, to better separate the EVR from ensuing voluntary recruitment and to further delineate the target-selective nature of the expected EVR strengthening with SAS. Finally, we also examined EVRs and RTs on a simple reaction task where a movement could be fully prepared prior to stimulus emergence, enabling comparison to results from the choice reaction task.

## Materials and methods

### Ethical Approval

A total of 17 subjects (10 females, 7 males; mean age: 22.6 years SD: 5.7) participated in these experiments. Subjects were volunteers who were mainly undergraduate students recruited by word of mouth. Two of the 17 subjects are the lead authors of this manuscript, and we observed no evidence that their results differed from those naive to the experimental goals. All subjects provided informed written consent, were paid for their participation, and were free to withdraw at any time. All but 3 subjects were right-handed, and all subjects had normal or corrected-to-normal vision, with no current visual, neurological, or musculoskeletal disorders. All procedures were approved by the Health Science Research Ethics Board the University of Western Ontario (HSREB 103341) and conformed to the Declaration of Helsinki.

### Apparatus and experimental design

Subjects were seated and performed reaching movements with their right arm in a KINARM End-point lab, moving the end-point of a robotic manipulandum in response to the appearance of visual stimuli that were occasionally accompanied by a loud auditory stimulus. Visual stimuli were computer-generated images produced by a projector (PROPixx project by VPixx, Saint-Bruno, QC, Canada) integrated into the KINARM setup, and projected onto an upward facing mirror. A shield below the mirror occluded direct vision of the hand, and hand position was represented by a real-time cursor (1 cm radius) projected onto the screen. Subjects were instructed to generate arm movements as quickly and as accurately as possible in response to stimulus emergence in an emerging target task (Kozak *et al*., 2020), moving either toward (a pro-reach) or away from (an anti-reach) the stimulus depending on an instructive cue provided at the start of each trial (see below). To ensure that all kinematic and electromyographic (EMG) data are aligned to the exact time of stimulus emergence and to control for possible delays in stimulus presentation by the projector, the precise time of stimulus emergence below the barrier was synchronous with the presentation of an accessory visual stimulus below a photodiode. This accessory stimulus was not seen by the participant, and photodiode output was fed to the KINARM platform. All kinematic and EMG data were aligned to photodiode onset. Throughout the entire experiment, a constant load of 2 Nm towards the participant and 5 Nm to the right was applied through the manipulandum in order to increase the activity of the right pectoralis muscle, so that the activity of this muscle would increase or decrease, respectively, following stimulus presentation in the preferred or non-preferred direction of the muscle. The same load was applied for all participants.

On a subset of trials, a loud acoustic stimulus was presented at the same time as the emergence of a visual target. The acoustic stimulus consisted of a 40 ms white noise burst delivered at an intensity of between 119 and 120 dB. A bilateral sound file was played through a digital output channel in the Kinarm setup and fed into a Rolls stereo line mixer/headphone amplifier, (model RM219) and then delivered bilaterally to Beyerdynamic CT 240 Pro headphones worn by the subject. This output was also routed to an analog in-channel on the KINARM platform, allowing us to confirm the synchronization of the auditory stimulus with visual stimulus emergence measured by the photodiode. Prior to the experiment, the sound intensity from each earpiece was calibrated by placing the earpiece on top of a GRAS Ear Simulator (model RA0039) with a 1 ⁄ 2 ″ microphone and held in place by 500g weight. Sound files were recorded with an M-Audio Fast Track Ultra audio interface and analyzed in Praat analysis software (Boersma, 2001). The sound intensity produced by the right and left earpiece was measured at 119.6 dB and 119.1 dB, respectively.

Subjects performed a number of variants of this task in different blocks of trials, and we will describe the results from two such blocks. The order of the blocks was randomized across subjects. Both blocks were variants of the emerging target task (Kozak *et al*., 2020), which increases the probability of observing EVRs on upper limb muscles (Contemori *et al*., 2021b, 2021a; Kozak & Corneil, 2021; Kearsley *et al*., 2022). The structure of this paradigm is provided in Fig. 1. Trials were separated by a 1.5s inter-trial interval. At the start of each trial, the configuration shown at the top of Fig. 1 was presented, with a barrier colored either red or green. The color of the barrier instructed the subject to prepare to make a pro- (toward) or anti- (away from) reach, relative to the side of stimulus emergence below the barrier. Subjects moved the cursor (1 cm radius) representing their hand position into a start location (1 cm radius), at which point a visual stimulus (1 cm radius) was placed above a barrier. After a 1000 ms hold period, during which subjects were required to maintain the 1 cm radius hand position cursor over the 1 cm radius start location (if not, the trial was reset; the tolerance was such that any portion of the hand position cursor had to touch the start location), the stimulus was depicted to travel as if it was following down an inverted “y” path at a speed of 15 cm/s for 500 ms before disappearing behind the barrier. The paradigm emulates a scenario where the junction of the y was obscured by a barrier, hence the stimulus appears to first disappear behind the barrier, and then emerge from beneath the barrier at either the right or left outlet. The outlets were approximately 20 cm lateral to and slightly above the starting position of the hand. Stimulus emergence was timed as if it was moving at a constant velocity behind the barrier, thus it appears to the participant that the stimulus was obscured behind the barrier for a fixed period of 500 ms on all trials. During the time the stimulus appeared to be behind the barrier, subjects were instructed to keep their hand at the start location, and to fixate a small notch at the bottom of the barrier (eye movements were not measured). At the time of what appears to be stimulus emergence, the stimulus was drawn as a full circle that continued to move along the inverted y path, and hence moved obliquely toward/lateral relative to participant midline. Upon stimulus emergence, subjects were instructed to respond as quickly and as accurately as possible and move to intercept the target on pro-trials with a 2-dimensional movement of the manipulandum, or move in the diametrically opposite direction on anti-trials. The trial ended if the hand cursor made contact with the stimulus on pro-trials, reached the diametrically opposite location on anti-trials, or if the stimulus moved off screen. On 25% of all trials, stimulus emergence was accompanied by a non-directional SAS.

**Figure 1.**
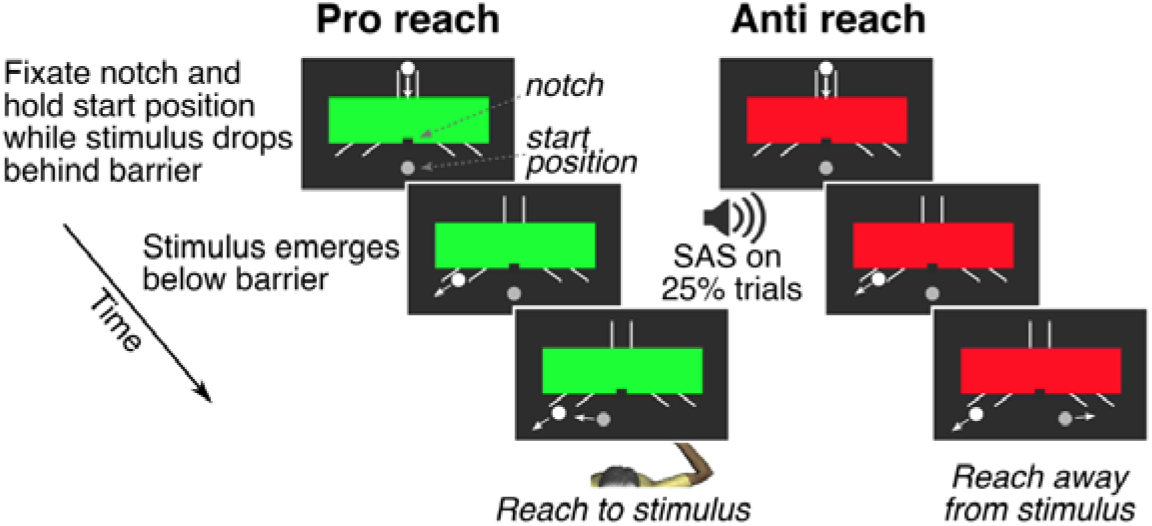
Behavioral paradigm. At the start of each trial, participants acquired the central start position with their hand (grey circle), and fixated a small notch at the bottom of the barrier. The barrier color conveyed the instruction to reach toward (green barrier, a pro-reach) or away from (red barrier, an anti-reach) the stimulus (white circle) upon its emergence below the barrier. On 25% of trials, a starting acoustic stimulus (SAS; 119-120 dB) was presented at the time of stimulus emergence. In a block of Choice reaction task trials, the stimulus could emerge at either the left or right outlet with equal probability. In a block of Simple reaction task trials, the stimulus only emerged at the left outlet.

In a block of Choice reaction task trials, the stimulus could emerge either to the left or right, and subjects were instructed to respond with either a pro-reach toward the stimulus (green barrier) or an anti-reach away from the stimulus (red barrier). Thus, there were 8 unique trial conditions: stimuli to the left or right, requiring a pro- or anti-reach, with or without a SAS. Subjects completed 1 block of 240 pseudorandomized trials. 60 (25%) trials contained a SAS, and 180 (75%) of trials had no SAS. Thus, there were 15 or 45 unique repeats of trials with or without a SAS, respectively.

In a block of Simple reaction task trials, the stimulus always appeared to the left, and subjects were instructed to either respond with a pro- or anti-reach. Subjects were explicitly informed of the left-sided stimulus presentation in this block, and they were told that this resulted in 100% certainty of whether a pro-reach to the left or an anti-reach to the right would be required at stimulus emergence. This task thus allowed for full preparation of the requested leftward or rightward hand movement. There were 4 unique trial conditions: a leftward stimulus requiring either a pro- or anti-reach, with or without a SAS. Subjects completed 1 block of 120 pseudorandomized trials, 30 (25%) or 90 (75%) of which contained a SAS or not, respectively. Thus, there were 15 or 45 unique repeats of trials with or without a SAS, respectively.

### Data acquisition and analysis

Surface electromyographic (EMG) recordings were made from the following targets: the clavicular head of the right pectoralis major muscle, the sternal head of the right pectoralis major muscle and right and left sternocleidomastoid (SCM) muscles. In all cases, recordings were made with double-differential surface electrodes (Delsys Inc., Bagnoli-8 system, Boston, MA). We found that the recordings from the clavicular and sternal heads of pectoralis major were essentially equivalent, so report the results from the clavicular head. EMG signals were sampled at 1 kHz, amplified by 1000, full-wave rectified off-line, and smoothed with a 7-point smoothing function.

Kinematic data were sampled at 1 kHz by the KINARM platform. RTs were detected based on acceleration and velocity criteria. For a given trial, we first found the point in time where the arm exceeded 10% of its tangential peak velocity. We then searched back in time for the latest point relative to stimulus presentation where the arm’s acceleration fell within a 99% confidence interval of arm accelerations when the arm was supposed to be stable. This 99% confidence interval was determined from all trials from the given subject based on the arm’s minor accelerations during a timeframe spanning from 100 ms before to 50 ms after stimulus appearance. Trials with RTs below 80 ms were excluded as anticipatory, which is supported by an analysis in the Choice reaction task showing that Pro-reach trials with RTs greater than this value were correct more than 80% of the time, whereas those started earlier were not. Trials with RTs exceeding 600 ms were excluded due to presumed inattentiveness. Overall, a total of 3.38% of trials were excluded in the Choice reaction task using the RT cutoffs, with the vast majority of being anticipatory movements. We applied the same RT criteria to data from the Simple reaction task, and rejected 34.4% off all trials, with virtually all such exclusions being anticipatory movements. All trials were also inspected by an analyst in a graphical user interface, which permitted rejection of trials with clearly anomalous movement sequences. Such rejected trials included those where the subject did not respond, where the limb was moving well before the stimulus appeared below the barrier, where the participant failed to reach the goal by moving less than half of the way toward the correct location, or produced multi-component movement sequences composed of three or more components. 3.1 ± 2.3% (mean ± SD) of all trials were rejected by the analyst for these reasons.

We retained movement sequences where subjects first moved in the wrong direction before correcting the reach to attain the goal. These movement sequences were termed wrong-way errors, and were more prevalent on anti- vs pro-reach trials (see Results). For such trials we determined the onset latencies in the incorrect as well as the correct directions. The former was determined as explained above, whereas the latter was determined as the time when the reach started to proceed in the correct direction. As detailed in the Results, for some analyses of EMG activity on anti-reach trials, we restricted analyses to those trials where subjects either moved directly away from the emerging stimulus, or moved no more than 50% of the distance toward the emerging stimulus, relative to where they landed on pro-reach trials, before correctly reversing the reach in the opposite direction. Our rationale here is that such mid-flight reversals indicate that subjects had consolidated the anti-reach instruction. We note that this 50% cutoff is arbitrary, and to satisfy ourselves that our results and conclusions were not due to this particular value, we re-ran all analyses after changing this cutoff to 25% (which excludes more anti-reach trials) or 75% (which excludes fewer anti-reach trials). In both cases, the qualitative nature of the results presented below, particularly regarding the latency and magnitude of the EVR on anti-reach trials, remained the same regardless of which cutoff was used.

As described previously (Corneil *et al*., 2004), we used a time-series receiver-operating characteristic (ROC) analysis to determine the presence and latency of the EVR in the Choice reaction task. Briefly, we conducted an ROC analysis for each point in time from 100 ms before to 300 ms after stimulus presentation. For each point in time, the area under the ROC curve indicates the likelihood of discriminating the side of stimulus presentation based only on EMG activity alone; a value of 0.5 indicates chance performance, whereas a value of 1.0 indicates perfect discrimination. While our past work (Wood et al., 2015; Kozak et al., 2021) determined the presence or absence of an EVR by conducting separately time-series ROC curves for the shorter- and longer-than-average RT subsets, this was not practical in the current dataset given the fewer number of repeats of each unique stimulus condition, and the relatively small variance in RTs. Instead, we found the time at which the slope of the time-series ROC changed by using the matlab function *ischange*; if this time fell within 70 and 120 ms, then we determined that an EVR was present, and the time at which the slope changed was determined to be the EVR latency. EMG magnitude in the EVR time window was calculated as the mean activity over the 80-120 ms interval post stimulus onset. Following subtraction of baseline activity, defined as the 500ms of activity prior to stimulus onset, these EMG magnitudes were normalized with respect to the maximum value of the ensemble-averaged PEC activity on left pro-reach trials without an SAS. Note that these EMG magnitudes were determined regardless of whether an EVR was identified. We note that this time-series ROC analysis is not possible in the Simple reaction task, since the stimulus is always presented to the left. While there are alternative methods for EVR detection that could have been used (Contemori *et al*., 2022; Kearsley *et al*., 2022), for the sake of simplicity we do not calculate EVR latencies for the Simple reaction task, and quantify EMG recruitment during the predefined interval of 80-120 ms after stimulus presentation.

### Statistical analysis

Unless otherwise stated, linear mixed models were used to investigate main effects and interactions. Linear mixed models were chosen over repeated-measures analysis of variance (ANOVA) because unlike ANOVAs, linear mixed models do not use list-wise deletion in the case of missing data points, allowing us to maximize the power and reduce the bias of our analysis. This applies where a participant may exhibit an EVR in one condition but not another (e.g., on trials with or without a SAS). The Satterthwaite method was applied to estimate degrees of freedom and generate p-values for the mixed model analyses. We investigated the effect of stimulus presentation side (left vs right), instruction (pro-reach vs anti-reach) and startle (no-SAS vs SAS), specifying these as fixed effects and participant ID as a random effect in the linear mixed models. Post hoc orthogonal contrasts with the Bonferroni correction method for multiple comparisons were used to investigate significant interactions between predictor variables, with an alpha of 0.05. We used paired t-tests to determine whether the SAS influenced EMG activity in an interval preceding the EVR, and to compare RT and EVR magnitude on trials based on Startle activity. We used a linear regression to correlate EVR magnitude versus RT across our sample. Data processing was done in MATLAB (R2021a), and statistical analyses were performed using jamovi (version 2.3, 2022), and MATLAB (R2021a).

## Results

### Choice reaction task - performance and movement RTs

Following trial exclusion, we retained a total of 3708 trials (90.9 ± 3.8%; mean ± SD) for further analysis (see Methods for exclusion criteria and frequency of different exclusion types). ‘Wrong-way’ error rates and RTs for each of the experimental conditions are displayed in Figure 2A and 2B, respectively. Participants made more mistakes on anti-reach trials than pro reach trials (15.2 ± 6.6% vs 3.4 ± 3.5% of trials, respectively) resulting in a main effect of instruction (*instruction*; β = 0.117, p = 1.43e-15, 95% CI [0.0926, 0.1422]). Participants also made more wrong-way errors on SAS than non-SAS trials (13.3 ± 7.5% vs 5.3 ± 3.1% respectively; *SAS*, β = 0.080, p =4.68e −9, 95% CI [0.0555, 0.1051]), which depended on the instruction given (*instruction x SAS*; β = 0.125, p =2.86e-6, 95% CI [0.0751, 0.1742]). A post hoc comparison showed that in anti-reach trials there were more wrong-way errors with SAS than without (22.3± 11.4% vs 8.0± 4.5%; p = 8.37e-12), but this was not the case with pro-reach trials (4.3 ± 5.4% vs 2.5± 2.7%; p = 1.000). There was no evidence that these results differed significantly as a function of the side of the target appearance (p>0.435 for all main or interaction effects involving *side*).

**Figure 2.**
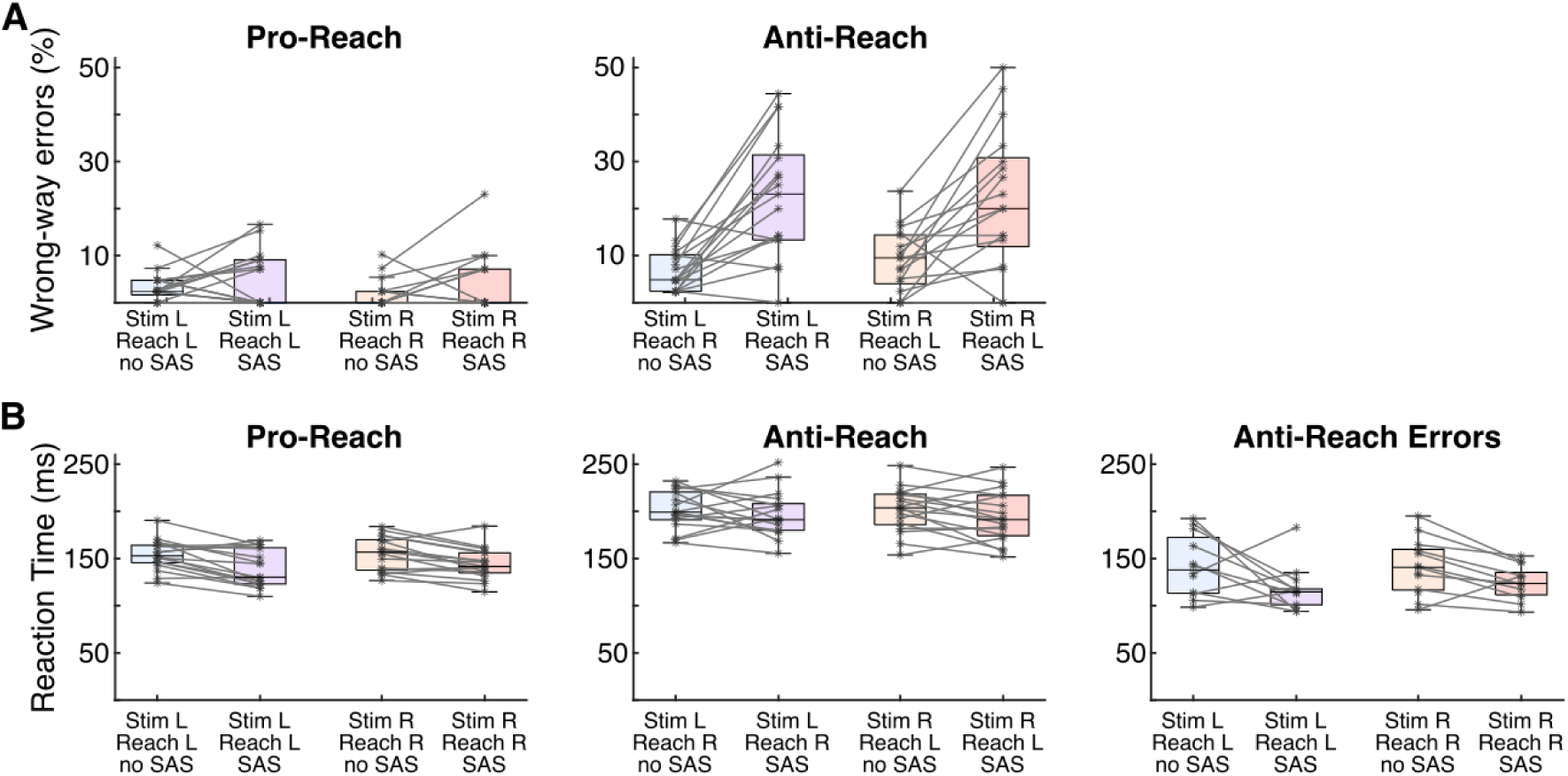
Behavioral results from Choice reaction task. Depiction of error rates (**A**) and RTs (**B**), for all 17 participants. Errors are defined as anti-reach trials where participants initially moved incorrectly toward the emerging stimulus, and then corrected the movement in mid-flight to reach in the opposite direction. In all cases, x-axis labels provide the response the subjects should have generated. For the RTs of anti-reach trials shown in B, the middle panel shows the RTs for the correct movement away from the simulus, whereas the right panel shows the RTs for the incorrect movement toward a stimulus on error trials. A given subject had to generate at least 2 such errors to be included in this panel. For boxplots, the *black, horizontal line* depicts the median across the sample, the *coloured portion* spans the 25th to 75th percentile, the *error bars* depict the span of data not considered outliers, the *asterisks* depict the mean of the observations from individual subjects, and the faint gray lines connect data from a given subject across trials with and without a SAS, where both values are available.

Movement onset latencies were shorter in pro- than in anti-reach trials (148±15 ms vs 198±18ms, respectively; *instruction*, β = 50.239, p= 5.28e-39, 95% CI [45.35, 55.13]). Note that for the wrong-way trials for anti-reaches, we included the onset latency of the movement away from the target (i.e. the instructed direction). The SAS significantly shortened movement onset latencies by, on average, 12 ms in pro-reach trials (142±16 vs 154±16 ms without SAS) and by 6 ms in anti-reach trials (195±20 vs 201±21 ms without SAS; *SAS*, β = −9.443, p = 2.50e-4, 95% CI [−14.34, −4.55]). There was no evidence for an interaction effect between the effect of the SAS and instruction (*instruction x SAS*, β = 6.4505, p =0.199, 95% CI [ −3.33, 16.23]), and no evidence for a main or interaction effect involving the side of target appearance (p > 0.3 for main or interaction effects involving *side*).

Across participants, the latencies of wrong-way movements (i.e. the RT of the movement towards the target) in anti-reach trials were shorter for SAS than non-SAS trials (122±20 and 146±31 ms, respectively; *SAS,* β =−21.21, p =0.002, 95% CI [−34.9, −9.48]) with no evidence for an effect of target side (*side*, β =2.55, p =0.701, 95% CI [−10.4, 15.46]). The maximum hand displacement in the wrong direction did not significantly differ between SAS and non-SAS trials (11.9±6.2% vs 10.7±5.3% of the distance to target; *SAS,* β =1.135, p =0.380, 95% CI [−1.65, 4.375]) but was significantly larger for wrong way movements to the left (13.5± 6.1%, relative to movement amplitude on pro-reach trials) than right (9.5 ± 5.1%; *side*, β =−3.55, p =0.030, 95% CI [−6.62, −0.477]).

### Choice reaction task – Effects of SAS on EVR Latency and response magnitude

Figure 3a-d shows the EMG responses in the pectoralis (PEC) muscle of a representative subject for each of the reaching conditions with and without a SAS. As the characteristic feature of the EVR, a band of increased PEC activity can be seen in the trials where the stimulus was presented on the left side (i.e left column) at 80-120 ms post stimulus onset, whereas in trials where the stimulus was presented at the right side (i.e. right column) a decrease in activity occurs in this time window. In pro-reaches (i.e. top row) this contrast in PEC activity between left and right stimulus presentation is more pronounced than in anti-reaches (i.e. bottom row). Figures 3b and 3d show the respective time-series ROC analyses for identifying the presence and latency of the EVR (see methods).

**Figure 3.**
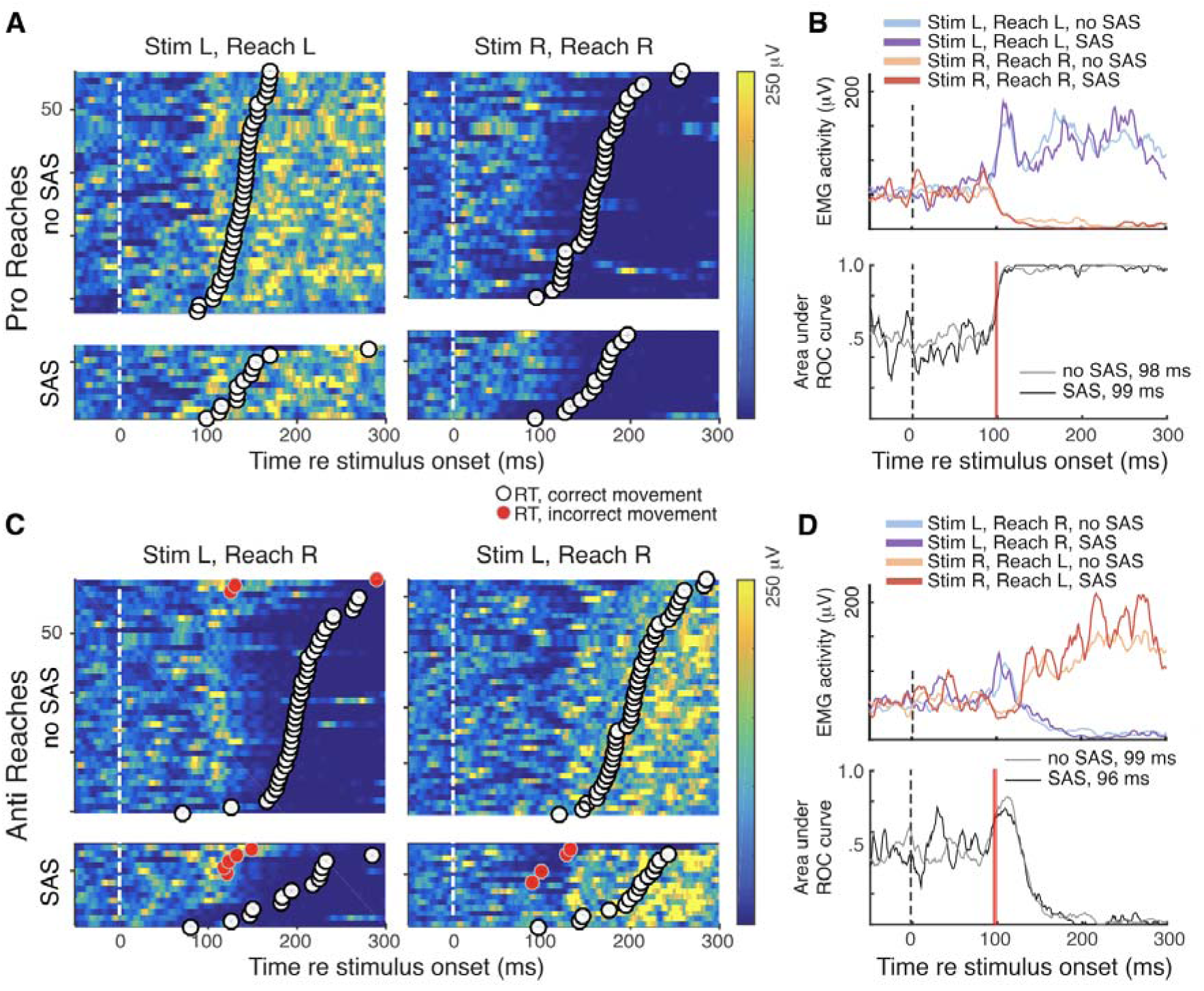
Representative EMG activity from the pectoralis muscle of an exemplar participant from Choice reaction task. EMG activity is shown in trial-by-trial heatmaps for pro-reach (**A**) and anti-reach trials (**C**). In each heat map, color conveys the magnitude of EMG activity aligned to stimulus emergence, with each row depicting an individual trial with trials ordered by the RT of the movement in the correct direction (white circles). Red circles on rows for anti-reach trials depict the RT of a wrong-way movement toward the emerging stimulus, which preceded the onset of the correctly-directed reach. Separate heat maps are depicted from trials with or without a SAS. **B, D**: Lines in the upper subplots depict the mean EMG activity for the four trial types. Lower subplots depict time-series ROC, calculated separately for trials with or without a SAS. Vertical red lines depict the time at which a change in time-series ROC was detected (values provided in each subplot).

All participants had a significant EMG discrimination time in the EVR window (70-120 ms), indicating the presence of an EVR in at least one condition. In pro-reach conditions we observed significant discrimination times in 12/17 participants with the SAS present and 16/17 participants in non-SAS conditions. 12/17 participants had a significant discrimination time in the non-SAS anti-reach condition, and 11/17 in SAS anti-reach condition. The Linear Mixed Model yielded no main effect of SAS on EVR latency (*SAS*, β = 2.25, p = 0.120, 95% CI [−0.526, 5.02]). Note that this model did not include *side* because to evaluate the EVR, right reaches are already compared to left reaches to determine the ROC curve and subsequently the EVR timing. Discrimination times (Fig. 4A) in pro-reaches (89±6ms) were slightly but significantly shorter than in anti-reaches (93±7 ms; *instruction*, β = 3.87, p = 0.010, 95% CI [1.085, 6.65]), irrespective of the SAS (*SAS x instruction,* β = 1.28, p = 0.645, 95% CI [−4.131, 6.70]). This small effect seems to be driven by the smaller-magnitude EVR on anti-reach trials, as well as by differences in the EVR detection using the change of slope detection method (see methods), as the method appears to be less sensitive in anti-reaches when the time-series ROC briefly increases before decreasing.

**Figure 4.**
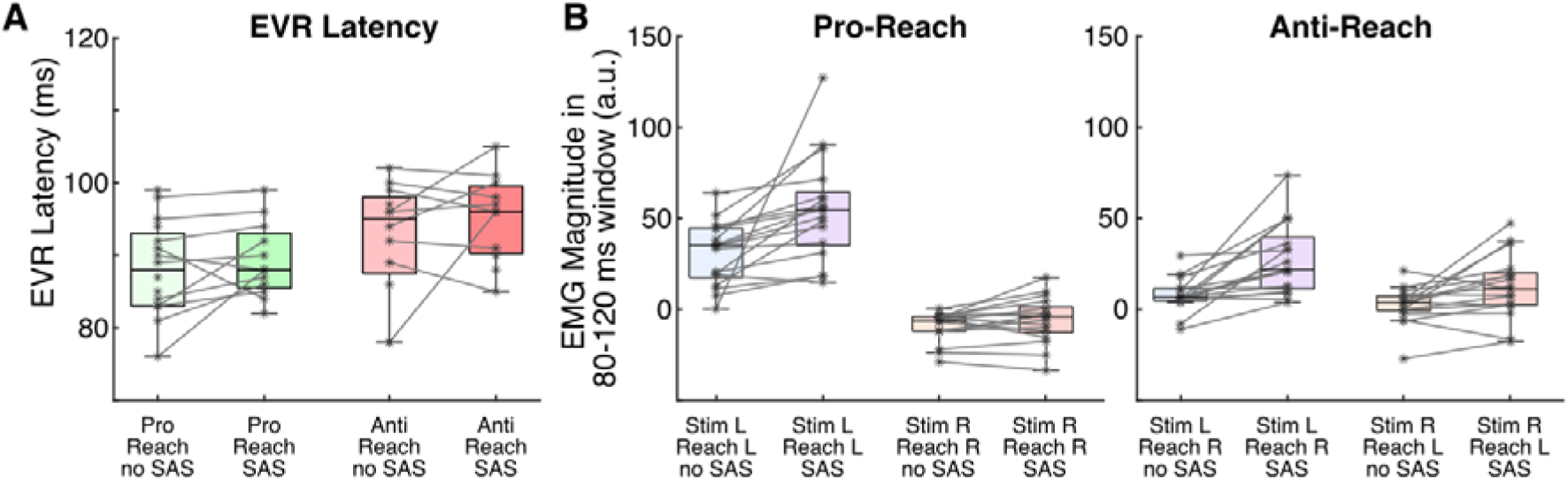
Quantification of the EVR in the Choice reaction task (Task A). Depiction of the latency (**A**) and magnitude (**B**) of the EVR for the 17 subjects in the sample. Same format as FIg. 2. Recall that EMG activity initially decreases following rightward stimulus presentation, which is why values may fall below zero (horizontal dashed line) in **B**.

Figure 4B shows the magnitude of PEC recruitment during the EVR window (80-120 ms), normalized relative to the maximum level of PEC recruitment aligned to reach onset averaged across all non-SAS left pro-reach trials. As expected for the EVR, PEC activity was significantly larger when targets were presented to the left than to the right (*side*, β = −30.32, p =4.37e-22, 95% CI [−35.22, −25.41]), and more so in pro- than anti-reaches (*side x instruction*, β = 39.73, p=1.75e-12, 95% CI [29.91, 49.54]). PEC activity was significantly larger with a SAS than without SAS (*SAS*, β = 14.38, p=8.07e-8, 95% CI [9.47, 19.28]). This effect of the SAS depended on the side of target presentation (*SAS x side*, β = −14.96, p =0.003, 95% CI [−24.77, −5.144]), but not on instruction (*SAS x instruction*; β = 1.00, p = 0.842, 95% CI [−8.81, 10.8161]). Post hoc analyses revealed that the SAS significantly increased PEC recruitment in leftward targets (p =6.57e-8) but not in rightward targets (p = 0.323).

### Relating EVR magnitude to movement RTs across SAS and non-SAS trials

Our task design in the Choice reaction task ensured that participants knew to generate a pro- or anti-reach on a given trial, but remained uncertain about whether the stimulus would emerge to the right or left. Despite this, participants generated pro-reaches with very short RTs (on average 142 ms or 154 ms with or without a SAS, respectively). When taking into account the electromechanical delay between the EMG signal and reach onset, this indicates that the forces arising from muscle recruitment during the EVR interval contributed to movement initiation. Across our sample, a SAS on pro-reach trials lowered RTs by 12 ms on average, ranging from a maximum reduction of 37 ms (168 or 131 ms on trials without or with a SAS) to a reduction of −3 ms (152 or 155 ms on trials without or with a SAS). Pro-reach data from these two subjects, along with the time-series ROC analyses, are shown in Fig. 5A and B. The subject in Fig. 5B with the smallest RT reduction had an EVR on pro-reach trials with or without a SAS and, as in the representative subject (Fig. 3), the SAS strengthened the magnitude of the EVR without changing its timing. In contrast, the subject with the largest RT reduction (Fig. 5A) is the only subject that did not have an EVR on pro-reach trials without a SAS. In this subject, the SAS produced a very prominent EVR, the timing of which resembled that observed in the rest of our sample. Thus, EVRs remained the earliest detectable change in muscle recruitment that depended on the side of stimulus emergence in the Choice reaction task.

**Figure 5.**
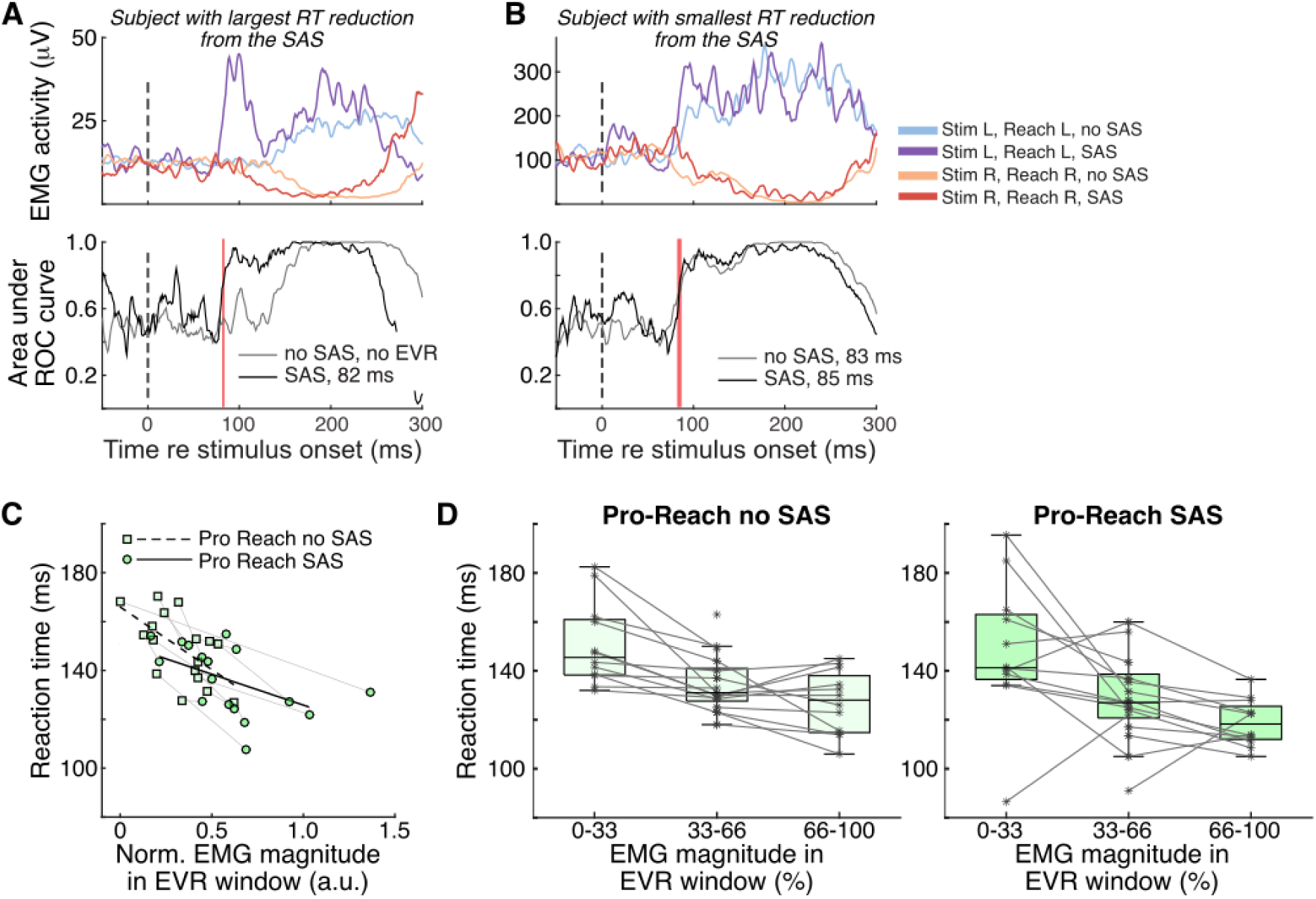
Negative relationship between RT and EVR magnitude. **A, B.** Mean EMG (top row) and time-series ROC (bottom row) for subjects where the SAS elicited either the largest (**A**) or smallest (**B**) reduction in RTs on Pro-reach trials, showing that a SAS provoked an EVR in both cases. Same format as Fig. 3B,D. **C.** Negative correlation between reaction time plotted as a function of normalized recruitment in the EVR window, for all 17 subjects for Pro-reaches in the Choice reaction task. Each symbol denotes the mean observation from a subject, with thin gray lines connecting observations with and without a SAS. Dashed or solid black line shows a linear regression for pro-reach trials without (r = −0.54, p = 0.026) or with (r = −0.55, p = 0.021) a SAS, respectively. **D.** Reaction time plotted as a function of binned EVR magnitude, for pro-left reach trials without (left subplot) or with (right subplot) a SAS. For all subjects, we derived the median RT associated with the normalized EVR magnitude within 3 equal sized bins of EVR magnitude. Same format as Fig. 2.

Prior research has established a negative correlation between EMG recruitment in the EVR interval and the RT on pro-reach trials (Pruszynski *et al*., 2010; Gu *et al*., 2016). These considerations lead us to question the degree to which the shortened RTs on SAS trials were associated with concomitant increases in EVR magnitude. Our hypothesis of intersensory facilitation of the SAS and a visual signal relayed through subcortical circuits predicts that RTs and EVR magnitudes should be related by a uniform relationship, with SAS trials leading to shorter RTs on average simply because of larger EVRs. To put it another way, a trial with a given magnitude EVR should have the same RT, regardless of whether a SAS was presented or not.

We addressed this question in a number of ways. First, we conducted an across-participant analysis where we plotted the mean magnitude of normalized muscle recruitment during the EVR interval as a function of mean reaction time, doing so separately for trials with or without a SAS. As shown in Fig. 5C, this revealed the expected normalized relationship, with the EVR magnitude being negatively correlated with the RT for pro-reach trials with (r = −0.60, p = 0.010) or without a SAS (r = −0.53, p = 0.028). Although the slope of these negative correlations were shallower for trials with a SAS, such a difference may be due to a basement effect where RTs could not go lower on SAS trials despite a few examples of large magnitude EVRs. Second, we conducted a within-participants analysis for leftward pro-reach trials, comparing the RTs on SAS and non-SAS trials that are matched for EVR magnitudes. For each participant, we binned the trials with respect to EVR magnitude (3 bins, bin width = 33%). Providing that there were sufficient SAS and non-SAS trials in a given bin (at least n = 1 of both), we derived the median RT for SAS and non-SAS trials in that bin. We then used a Wilcoxon signed-rank test to RTs across participants and bins (Fig. 5). RTs became faster with greater EVR magnitudes, but there were no significant differences between SAS and non-SAS trials (adjusted alpha = 0.05/3 = 0.0167; Bin 0-33, p = 0.622; Bin 34-66, p = 0.058; Bin 67-100, p = 0.097).

### Generalized startle reflex activity in upper limb and neck muscles that precedes the EVR

While the finding of enhanced EVR magnitudes with SAS in leftward but not rightward targets (Fig. 4) argues against a generic effect on PEC recruitment in this time window of interest, we further explored whether the SAS elicited a reflexive startle response before the EVR. Here, we took advantage of our recordings not only from PEC, where activity is related to the reaching task, but also from our recordings of bilateral sternocleidomastoid (SCM). Although SCM recordings are commonly used to assess the presence or absence of startle reflexes during StartReact experiments (for review see (Carlsen & Maslovat, 2019)), the typical time interval of up to 120 ms after the SAS in such assessments overlaps with the EVR interval (80-120 ms after stimulus emergence); thus we cannot use traditional methods to assess the presence or absence of startle reflexes on a trial-by-trial basis. We explored the time course of averaged activity from PEC and bilateral SCM after stimulus emergence, pooling across pro- and anti-trials and side of stimulus emergence, but doing so separately for SAS and non-SAS trials. We normalized the average activity of these muscles to the activity in the 500 ms preceding stimulus presentation, and then subtracted the activity on non-SAS from SAS trials. This analysis produces a difference curve where any increase in EMG activity in the time after stimulus emergence is attributable to the presence of the SAS. As shown in Fig. 6, the presence of the SAS increased activity in two intervals, one soon after the SAS (starting at ∼20 ms for bilateral SCM, and ∼30 ms for PEC), and another later on in the EVR interval. On PEC, this latter response during the EVR interval is expected because of the asymmetric effect of the SAS, as it increases recruitment more following left stimulus than it decreases it following right stimulus emergence (Fig. 4). A similar pattern of recruitment is also apparent in L-SCM starting at around 100 ms, although such recruitment was less common than in PEC and was observed in only a few subjects. We therefore focus on the earlier change in muscle recruitment, as such activity evolved well before the EVR. To assess the significance of these results across our sample, we ran sample-wise signed-rank tests to identify where this excess activity was significantly different from 0 (p < 0.05) for at least 10 consecutive samples. In PEC we found significant SAS-induced activity for a brief interval between 30 and 50 ms after stimulus emergence, well before the EVR (Fig. 6A). In bilateral SCM, there was also brief and very early (starting at ∼25 ms) increased EMG activity in SAS trials (Fig. 6B, C).

**Figure 6.**
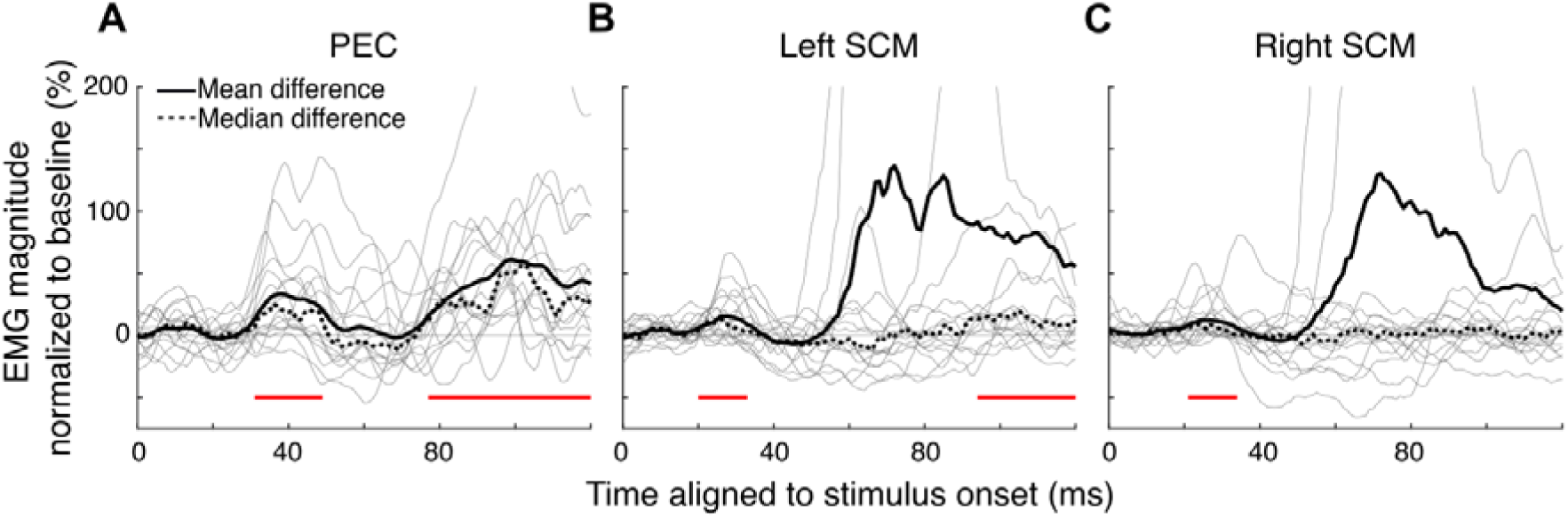
A SAS increases recruitment of right PEC and bilateral SCM activity before the EVR. Time-series of the difference in baseline-normalized EMG activity immediately after stimulus emergence due to the SAS, conducted separately for right-PEC (**A**), left-SCM (**B**), and right-SCM (**C**). In each subplot, thin lines show data from individual subjects, and red lines denote periods where significantly greater EMG activity was observed across our sample on SAS vs non-SAS trials.

We explored the trial-by-trial influence of such early recruitment on subsequent muscle recruitment and behaviour in a few ways. First, on each trial, we determined whether EMG recruitment in this early interval exceeded by two standard deviations the mean activity in a baseline interval determined from the 200 ms preceding stimulus emergence. Trials with or without significant activity are termed PEC+/PEC- or SCM+/SCM-trials, respectively, depending on which muscle is being assessed for this early startle activity. We ran this for all trials (regardless of whether a SAS was presented or not), and found that a SAS slightly but significantly increased the proportion of trials where significant activity was detected on either SCM muscle in an interval from 20 to 50 ms after stimulus emergence (SCM+ trials: 14.0 +/− 4.4% on trials without a SAS vs 20.0 +/− 7.3% of trials with a SAS; one-way paired t-test Bonferroni-corrected for multiple comparisons, p =4.80e-3, t(16) = −2.94), or on PEC muscle 30 to 60 ms after stimulus emergence (PEC+ trials: 44.4 +/− 7.0 % on trials without a SAS vs 52.2 +/− 14.1% of trials with a SAS; p = 0.017, t(16) = −2.33). On SAS trials, the presence of activity in this early interval did not relate to significantly shorter pro-reach reaction times using either SCM (Pro-reach RTs in either direction = 133.1 +/− 13.4 ms vs 130.6 +/− 16.6 ms on SCM- vs SCM+ trials; p = 0.171, t(15) = 0.983; excluding one subject within insufficient SCM+ trials) or PEC activity (Pro-reach RTs to the left = 130.6 +/− 19.3 ms vs 125.9 +/− 15.3 ms on PEC- vs PEC+ trials; p = 0.112, t(16) = 1.264). Consistent with this, the magnitude of PEC recruitment in the EVR interval on pro-reach trials to the left did not increase in the presence of significant startle activity for either SCM (relative to the EVR on SCM-trials, EVR on SCM+ trials = 1.27 +/− 1.06, p = 0.299, t(14) = 0.541; excluding two subjects with insufficient SCM+ or SCM-trials since only leftward trials were analyzed) or PEC (relative to the EVR on PEC-trials, EVR on PEC+ trials = 0.86 +/− 0.25, p = 0.999, t(16) = −3.618). Thus, although a SAS increased the recruitment of bilateral SCM and right PEC in a brief ∼30 ms interval that preceded the EVR, such recruitment had little influence on the magnitude of recruitment in the subsequent EVR interval, or on the ensuing reach reaction time.

### Simple reaction task - performance, movement RTs, and EVR magnitudes

The shortening of RTs in the presence of a SAS due to the StartReact effect is most commonly observed in experiments where subjects have foreknowledge of the requested response. In a separate block of trials, we therefore collected behavioral and EMG data from a Simple reaction task where stimuli always emerged to the left, to which participants responded with a left (pro-reach) or right (anti-reach) response, depending on the conveyed instruction. Compared to the Choice reaction task, we observed a large number of anticipatory responses (RTs < 80 ms; 36.9% vs 3.4% in Simple vs. Choice reaction task, respectively). Some subjects produced anticipatory responses more than half the time, hence we analyzed data only from the remaining 11 subjects that produced anticipatory responses on less than half of all trials.

We show data from a representative participant in Figure 7 (same participant as in Fig. 3). Behaviorally, the RTs on anti-trials are quite similar to those on pro-reach trials, and this participant did not generate wrong-way reaches toward the emerging stimulus on anti-reach trials (compare heatmaps and RTs in left columns of Figs. 3 and 7). Second, while prominent EMG recruitment during the EVR interval is apparent on pro-reach trials in the simple task (Fig. 7A,B), EMG recruitment during the EVR interval is absent on anti-reach trials (Fig. 7C,D). Thus, it appears that this participant fully prepared the motor program for the pro- or anti-reach before stimulus emergence. Finally, while the SAS further shortened RTs for both pro- and anti-reach trials, the SAS only augmented EMG activity during the EVR interval on pro-reach trials; we observed little to no increase in EMG activity in this interval following leftward stimulus emergence on anti-reach trials.

**Figure 7.**
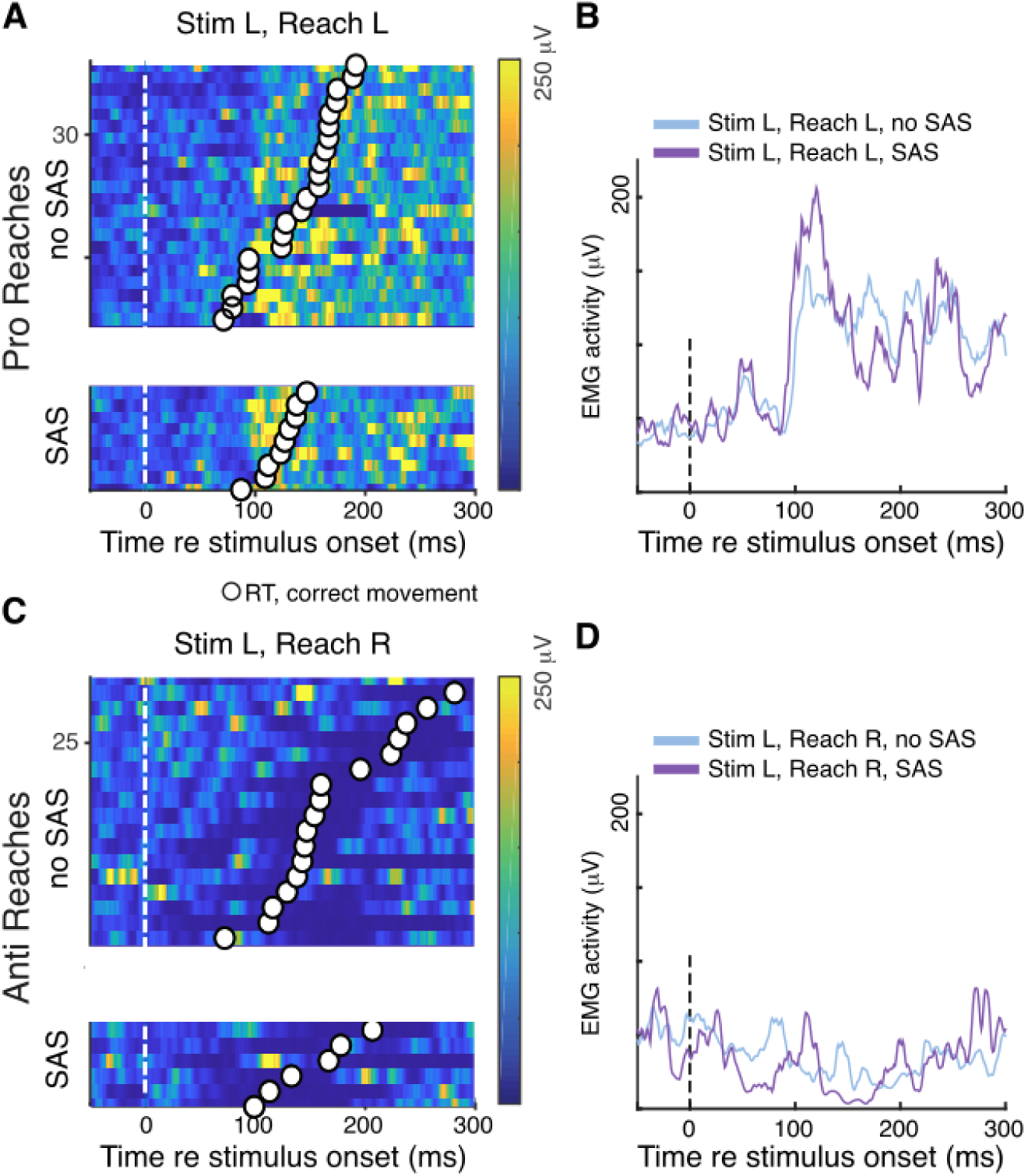
Representative EMG activity from the pectoralis muscle of an exemplar participant from Simple reaction task. Same participant and format as FIg. 3, excepting that a time-series ROC plot was not generated given the absence of trials with the stimulus emerging to the right.

We quantified the RTs and magnitude of EMG activity in the EVR interval across those 11 subjects that did not routinely anticipate stimulus emergence. The SAS significantly shortened RTs by 26 ms on average (Fig. 8A; 129± 21ms and 155±20ms for SAS and nonSAS respectively; SAS, β = −20.54, p = 3.76e-8, 95% CI [−27.05, −14.0]), irrespective of the instruction (*SAS x instruction*, β = 1.98, p = 0.767, 95% CI [−11.05, 15]) or the task (*SAS x task*, β = 10.58, p =0.116, 95% CI [−2.44, 23.6]). There was an interaction effect between task and instruction (*task x instruction*, β = 48.06,, p = 4.71e-10, 95% CI [35.04, 61.1]); a post hoc analysis showed that, in contrast to the Choice reaction task, we observed no significant difference between the RTs of pro- vs anti-reach trials in the Simple reaction task (Fig. 8A; 141±22 ms in pro- vs 142±19ms in anti-reaches; p = 1.000). Further, RTs for both pro- and anti-reaches in the Simple reaction task were comparable to the RTs for pro-reaches in the Choice task (p = 1.000), whereas the choice anti-reaches showed significantly different RTs (Fig. 8A, p = 3.73e-16).

**Figure 8.**
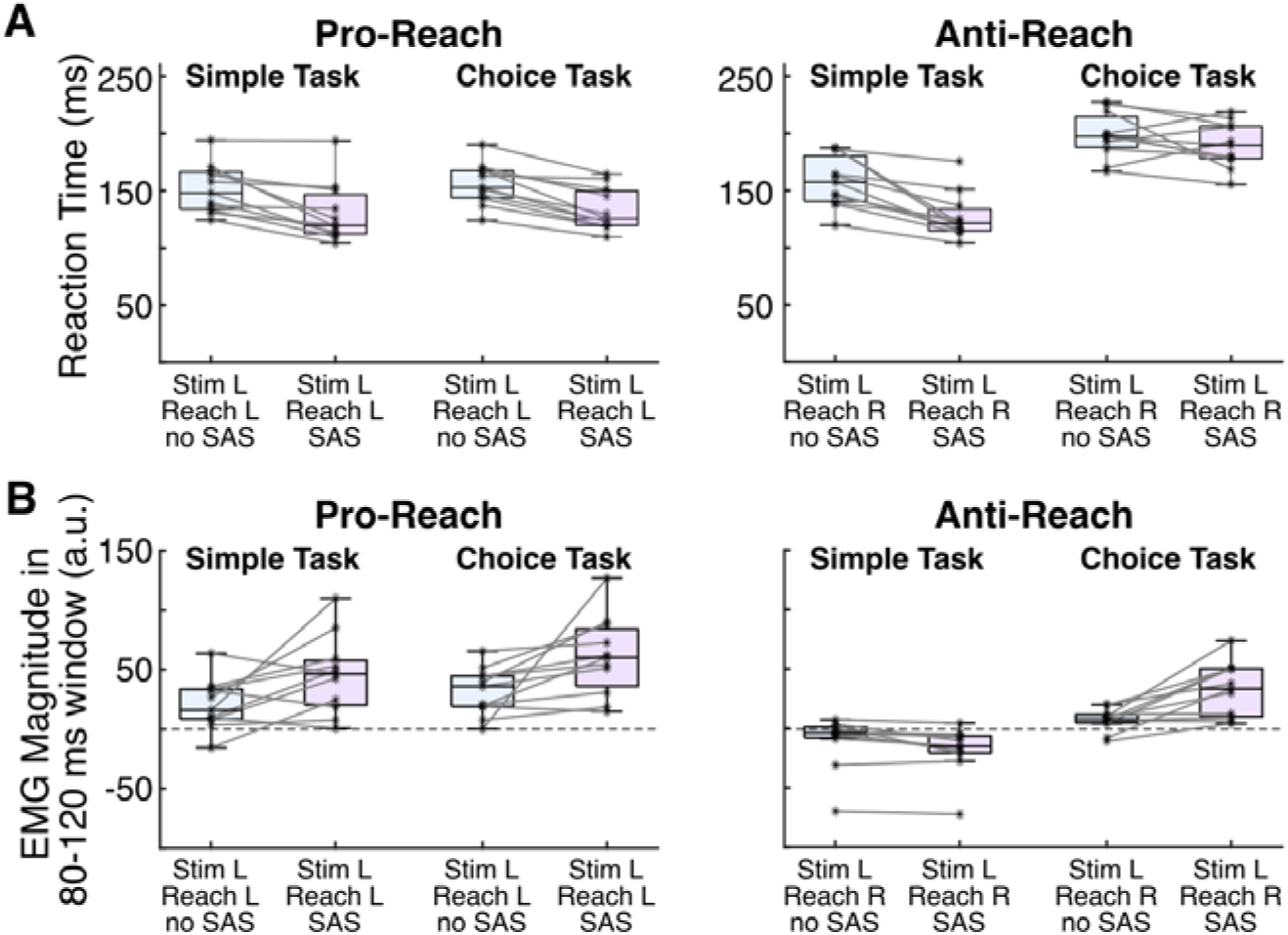
Quantification of behavior and EMG activity in Simple reaction task, compared to Choice reaction task. Depiction of RTs (**A**) and EMG magnitude in EVR interval (**B**), shown for the 11 subjects with sufficient data in the Simple reaction task (hence the subtle differences with Figs. 2 and 4). Same format as Fig 2.

In terms of EMG activity, PEC recruitment in the EVR interval across both tasks was significantly larger in pro- than in anti-reaches (*instruction*, β = −36.8, p = 4.34e-12, 95% CI [−45.49, −28.174]). Yet, this effect was dissimilar between tasks (*task x instruction;* β = 20.3, p = 0.025, 95% CI [2.97, 37.595]). While post-hoc tests showed that PEC recruitment was significantly different between pro- and anti-reaches in both tasks, the difference was greater in the Simple than the Choice reaction task (Choice; p = 3.58e-4, Simple; p = 8.38e-10). Post hoc tests revealed differences between almost all conditions, the only comparisons that did not show significant differences for the effect of task x instruction were simple and choice pro-reaches (p = 0.224), and simple pro-reach versus choice anti reach trials (p = 0.209). PEC recruitment was significantly larger in trials with SAS (*SAS*, β = 18.1, p = 1.10e-4, 95% CI [9.46, 26.769]) and during the Choice task (*task*, β = 23.4, p =1.30e-6, 95% CI [14.74, 32.050]), with a significant interaction (*task x SAS;* β = 18.7, p = 0.038, 95% CI [1.37, 35.994]). The SAS increased EVR magnitudes on pooled Choice task trial types (p= 2.32e-4), but its pooled effect across Simple task trial types was neutral (p = 0.988). Finally, there was an interaction effect between SAS and instruction (*SAS x instruction*; β = −17.8, p = 0.048, 95% CI [−35.12, −0.495]). The SAS had a potentiating effect on pro-reach trials when pooled across tasks (p = 2.98e-4), while its pooled effect in anti-reaches across both tasks was neutral (p = 0.870). While this neutral effect appears to be driven by opposing effects of the SAS in Simple vs. Choice task anti-reach trials (see figure 8B), the three-way interaction did not reach significance (*SAS x instruction x task*; β = 28.4, p = 0.113, 95% CI [−6.26, 62.997]). In sum, and in contrast to the Choice task, the effect of the SAS on anti-reaches in the Simple task (i.e., stimulus left, reach right) suppressed PEC recruitment during the EVR interval and produced RTs comparable to those observed on pro-reach trials.

## Discussion

We examined the effect of a SAS on behavior and upper limb muscle activity as human participants made pro- or anti-reaches in an Emerging Target task. The task promoted reactive RTs and the generation of short-latency bursts of muscle activity termed EVRs, even on trials without a SAS. In separate blocks of trials, the side of stimulus emergence could be varied (a Choice reaction task where responses could not be fully prepared) or be fixed (a Simple reaction task permitting full response preparation). The SAS lowered RTs in both tasks, and increased the magnitude of EVRs without altering its timing. Our results affirm that a SAS can reliably shorten RTs of reactive movements in select Choice reaction tasks. We surmise that the subcortical visuomotor pathway that produces EVRs is sufficiently primed prior to stimulus emergence in the Emerging Target task. In such scenarios, the hastening of RTs arises from intersensory facilitation within the reticular formation between the SAS and visually-derived signals relayed along a subcortical visual pathway; advanced preparation of a specific motor response and its release by the SAS, as in the StartReact effect, is not required.

Our RT results in a Choice reaction task (Fig. 2B) complement similar reports of how a SAS can shorten RTs of on-line lower limb corrections to displaced targets or obstacles (Reynolds & Day, 2007; Queralt *et al*., 2008), and demonstrate that the influence of the SAS can be observed for reactive movements of the upper limb initiated from a stable posture. Importantly, given that EVRs can also be observed on the lower limb (Billen *et al*., 2023), we suggest that past RT effects for on-line corrections of the lower limb may have arisen from strengthening rather than shortening signalling along a fast subcortical visuomotor pathway (Reynolds & Day, 2007). Given that a hastening effect of a SAS on RTs is generally not observed in Choice reaction tasks initiated from a stable posture (Carlsen *et al*., 2004, 2009; Forgaard *et al*., 2011; Maslovat *et al*., 2012; Marinovic *et al*., 2017)), what is distinct about the Emerging Target task? The Emerging Target task promotes a readiness to respond via strong top-down priming of a subcortical visual pathway due to implied motion and temporal certainty about the timing of stimulus emergence (Kozak *et al*., 2020; Contemori *et al*., 2021b). Consequently, pro-reach RTs with or without a SAS were essentially identical in both the Choice and Simple reaction tasks (Fig. 8). Similar facilitating effects of a SAS are also seen in launching interceptive actions (Tresilian & Plooy, 2006), and in promoting accurate responses in a forced RT paradigm (Heckman *et al*., 2023). All of these paradigms promote a degree of response urgency which may be an important factor in dictating reactive responses even without a SAS. As seen in the work by Heckman and colleagues (2023), a SAS in such scenarios can facilitate congruent movements directed toward a stimulus (pro-reaches in our case) or voluntary movements directed elsewhere (e.g., the RTs on correct anti-reach trials).

Is it possible that subjects in the choice reaction task prepared alternative motor programs in parallel in advance, which were then influenced, or perhaps even released, by the SAS? Evidence from multiple motor systems clearly shows that humans or primates can prepare a limited number of alternatives in advance that need not affect motor output (Basso & Wurtz, 1997; Dorris & Munoz, 1998; Cisek & Kalaska, 2005; Quoilin *et al*., 2019), so a systematic test of the influence of preparing multiple alternatives would require introducing more potential target locations. However, numerous considerations suggest that neither advanced preparation of parallel motor programs nor the SAS itself explains our results and those in the literature. First, while the phenomena of EVRs and SAS-induced effects on RT in reactive tasks benefit from the preparation of specific motor programs, such preparation is not critical; robust EVRs can be evoked even in the scenarios with up to twelve potential reach target (Selen *et al*., 2023) or when either limb could be used to reach to up to seven potential targets (Kearsley *et al*., 2022), and a SAS facilitates accurate responses in conditions of multiple potential targets in forced RT paradigm (Heckman *et al*., 2023). Our data also show that a SAS did influence neck and upper limb muscle activity within 20-40ms, which we attribute to a non-specific acoustic startle reflex (Brown *et al*., 1991b, 1991a). However, this phase of recruitment was not direction specific even on PEC; direction specificity only emerged later, i.e. during the EVR interval, and even then the timing of the EVR was not influenced by the SAS. This absence of SAS effects on EVR discrimination time is consistent with the findings of Glover and Baker (2019). In the EVR interval in the Choice reaction task, we also found that the SAS selectively increased PEC activity for leftward, but not rightward targets, regardless of whether participants were instructed to reach towards or away from the target. Thus, we saw no evidence of the SAS releasing a default motor program in the Choice reaction task. Our results speak to the SAS acting as an accessory stimulus that increases the excitation of the fast visuomotor network, such that it facilitates phases of muscle recruitment influenced by the emerging visual stimulus after the earliest startle reflexes. The fixed timing of the EVR reinforces our supposition that the pathway underlying the EVR represents the shortest pathway capable of transforming visual inputs into target-directed reaching actions (Gu *et al*., 2018; Contemori *et al*., 2023).

Our recordings of upper limb muscle activity demonstrate a consistent relationship between the earlier phase of stimulus-directed recruitment, the EVR, and subsequent RT. In the Choice reaction task, the SAS enhanced EVR magnitude but not timing. Such enhancement correlated with reduced RTs (Fig. 2B; Fig. 5C&D), and related to the increased propensity for wrong-way errors on anti-reach trials (Fig. 2A). Importantly, these lower RTs and increased wrong-way errors on anti-reaches were independent of target direction, which again speaks to the target-selective nature of EVR enhancement. These results affirm that the EVR, while brief in duration, leads to the production of relevant forces capable of initiating limb motion (Gu *et al*., 2016). Indeed, a subject-by-subject (Fig. 5C) and trial-by-trial comparison (Fig. 5D) of the relationship between EVR magnitude and RT shows that a given EVR relates well to a given RT, regardless of the presence or absence of a SAS. While there are undoubtedly non-linearities in how muscles generate force, in the context of the Choice reaction task experiment there appears a fairly straightforward explanation that the effect of the SAS on RTs is largely due to the production of a larger EVR. This was true even in the one subject (Fig. 5A) that did not produce an EVR on trials without a SAS, but had both large EVRs and the largest degree of RT shortening when a SAS was presented. Our interpretation from this example is that intersensory facilitation between the SAS and signaling from the visual stimulus was sufficiently strong to evoke an EVR in the periphery, whereas signaling from the visual stimulus alone was not.

We surmise that a true StartReact effect, wherein a SAS led to the involuntary ‘release’ of a prepared motor program, did occur in our Simple reaction task. Here, depending on instruction, left stimulus emergence requires a leftward pro-reach or a rightward anti-reach. Consistent with subjects preparing a specific motor program in advance of stimulus emergence, RTs on anti-reaches were ∼50ms faster than in the Choice reaction task, and as fast as those on pro-reaches. Furthermore, the strong recruitment in the EVR interval that was augmented by the SAS on pro-reach trials is completely absent on anti-reach trials, regardless of the presence or absence of the SAS (Figs. 7, 8). In the Simple reaction task, subjects had more than 2 seconds to consolidate the instruction to prepare for pro-reach responses to the left or anti-reach responses to the right, which apparently provides sufficient time on anti-reach trials to fully suppress the EVR to the leftward stimulus, which in this case acts as a signal to reach to the right. Such contextual suppression of the EVR resembles that observed in delayed reaching tasks (Pruszynski *et al*., 2010), and how EVRs from a given stimulus can be mapped onto different responses depending on task-relevant parameters (Gu *et al*., 2018; Contemori *et al*., 2023).

The reticular formation has been strongly implicated in both the StartReact effect (Valls-Solé *et al*., 1999; Nonnekes *et al*., 2015; Carlsen & Maslovat, 2019) and the phenomenon of EVRs (Corneil & Munoz, 2014; Contemori *et al*., 2023). Indeed, the reticular formation has the requisite connections to the motor periphery to detail the task-appropriate motor commands that are hastened by presence of a SAS in the case of the StartReact effect, or augmented in the case of intersensory facilitation. In the Choice reaction task used here, the reticular formation is a likely node for intersensory convergence between signals arising from the SAS and the emerging visual stimulus. Intersensory effects are also possible within the intermediate and deep layers of the superior colliculus, given its role in multisensory integration (Stein & Meredith, 1993) and inputs into startle circuitry (Fendt *et al*., 2001). Previous work examining multisensory integration in the SC of awake behaving monkeys has attributed the reductions in saccadic RT largely to changes in the timing and/or magnitude of saccade-related rather than visually-related signals (Frens & Van Opstal, 1998; Bell *et al*., 2005). However, such studies have used localizable acoustic stimuli with intensities <= 60 dB, hence the effect of a much louder SAS on visually-derived transients within the intermediate and deep layers of the primate SC is unknown.

There are a number of implications of our results. First, the magnitude of RT reduction alone cannot be used to differentiate behavioural effects due to a StartReact effect from intersensory facilitation. Tasks with a degree of response urgency, such as the one we used, engender shorter RTs on non-SAS trials to begin with, limiting the degree to which a SAS can further shorten the RTs of accurate movements. Indeed, the RT reductions we observed were similar in the Choice reaction task (∼12 ms) and the Simple reaction task (∼20 ms), both of which are in the range of reductions usually attributed to intersensory facilitation (Nickerson, 1973). However, the EMG results were consistent with intersensory facilitation for the Choice reaction task but a StartReact effect for the Simple reaction task. Second, EMG recordings from multiple muscles revealed that the SAS was sufficiently intense to provoke early, generic startle-related activity that was not dependent on the side of target presentation. The fact that such activity had little trial-by-trial influence on subsequent muscle recruitment in the EVR interval or on behaviour in the Choice reaction task is not what would have been expected of a StartReact mechanism (McInnes *et al*., 2021), but is consistent with intersensory facilitation and with previous results in the lower limb (Reynolds & Day, 2007). Third, such early startle-related activity was more prevalent on the pectoralis rather than the sternocleidomastoid muscle, despite the latter being the customary target for a trial-by-trial indicator of startle-based recruitment. Thus, there may be value in examining muscles in addition to, or perhaps other than, SCM depending on postural demands. Regardless, given the very rapid responses engendered by the emerging target task, the assessment window for startle-related recruitment necessarily had to be constrained to an interval preceding the EVR. Ultimately, future studies with a SAS in clinical or neurophysiological settings may benefit from incorporating paradigms that promote a degree of response urgency. Conversely, presentation of a SAS may increase the probability of observing EVRs in stroke patients, given the facilitating effect of a SAS on upper limb movements in this population (Honeycutt & Perreault, 2012; Honeycutt *et al*., 2015; Marinovic *et al*., 2016).

Taken together, our results provide compelling evidence that the observed RT shortening with SAS in the Choice task arise from intersensory facilitation of the fast visuomotor network, rather than a StartReact effect that invokes release of a partially or fully prepared motor program. EVR timing in the Choice task remained unaffected by a SAS, and enhanced PEC recruitment was selective to left-sided target presentation, indicating that lateralized PEC recruitment was not triggered by the SAS, but by the emerging visual target. A limitation of this study is that we did not record EMG from agonist muscles for rightward reaches (e.g. posterior deltoid). Yet, the behavioural results suggest that such recordings in the Choice reaction task would mirror those from PEC, given the similar overall RTs as well as the similar SAS effects on RTs and wrong-way errors between leftward and rightward targets. Given our supposition of intersensory facilitation being the underlying mechanism of the observed RT shortening with SAS, why then have previous reports largely failed to observe an influence of the SAS on RTs in Choice reaction tasks? A number of possible, and not mutually exclusive, explanations arise. First, a low level of response readiness in past tasks, perhaps due to the number of potential targets and/or uncertainty about the exact time of stimulus onset, engendered longer RTs which were generated after the SAS’ influence had dissipated. Second, in the context of reaching movements, it is possible that the SAS did facilitate small or subthreshold signaling along a fast subcortical visuomotor pathway, but such signaling was not sufficient to produce forces to overcome the arm’s inertia. Third, previous studies that did observe very fast RTs with SAS under Single but not Choice task conditions involved finger, wrist or elbow movements (Carlsen *et al*., 2004, 2009; Forgaard *et al*., 2011; Maslovat *et al*., 2012; Marinovic *et al*., 2017). As axial muscles are known to express stronger EVRs than distal muscles (Pruszynski *et al*., 2010), these movements may not equally benefit from SAS-induced facilitation of the fast visuomotor network. As such, and in agreement with the views expressed in recent review papers (Nonnekes *et al*., 2015; Marinovic & Tresilian, 2016; Carlsen & Maslovat, 2019), there is not a single unifying mechanism that explains how a startling acoustic stimulus expedites reactions times across all paradigms and effectors.

## ADDITIONAL INFORMATION

## Data availability statement

The authors confirm that the data supporting the findings of this study are available within the article. The raw data will be made available by the authors, without undue reservation.

## Competing interests

The authors declare no competing or conflicting interests.

## Author contributions

All experiments took place at the University of Western Ontario. VW and BDC contributed to study conception. VW, ALC, EAM, and BDC contributed to study design. VW and SLK collected the data and with BDC organized the database and performed data and statistical analyses. VW, SLK and BDC wrote the first draft of the manuscript. All authors contributed to manuscript revision, and read and approved the final version submitted for publication.

## Funding

VW was supported by a Netherlands Organisation for Scientific Research (NWO) Vidi grant (91717369) and an Erasmus+ Staff Mobility Grant. This work was supported by operating grants to BDC from the Natural Sciences and Engineering Research Council of Canada (NSERC) [RGPIN-311680, −04394-2021], and the Canadian Institutes of Health Research (CIHR) [MOP-93796, −142317; PJT-180279]. SLK was supported by Master’s and Doctoral scholarships from NSERC, and from Mitacs and the Parkinson Society of Southwestern Ontario. ALC was supported in part via an NSERC CREATE grant. The equipment used in this work was funded by the Canadian Foundation for Innovation.

## Notes

### Competing Interest Statement

The authors have declared no competing interest.

### Summary of Updates

Throughout our manuscript we are now very precise in our use of the term StartReact effect to refer to a mechanism, which is distinct from the behavioural effect of the hastening of RTs by a startling acoustic stimulus. This is important, as one of our major conclusions is that the hastening of RTs by a startling acoustic stimulus in the Choice reaction task is consistent with intersensory facilitation rather than the StartReact mechanism. We strive to clarify this throughout the entire manuscript. gWe have performed additional analyses to show that this interpretation holds across the range of RT benefits afforded by the startling acoustic stimulus. We have also examined those subjects with the largest (Fig. 5A) or smallest (Fig. 5B) RT gains afforded by the startling acoustic stimulus. The SAS bolstered the EVR in both subjects, and we now also show the inverse relationship between EVR magnitude and RT in Fig. 5C. In the Discussion, we provide further context to our conclusions for underlying neural mechanisms both in Choice reaction tasks with a high degree of urgency where intersensory facilitation effects predominate, and for past studies in the literature.

